# Comprehensive single-PCR 16S and 18S rRNA community analysis validated with mock communities and denoising algorithms

**DOI:** 10.1101/866731

**Authors:** Yi-Chun Yeh, Jesse C. McNichol, David M. Needham, Erin B. Fichot, Jed A. Fuhrman

## Abstract

Universal SSU rRNA primers allow comprehensive quantitative profiling of natural communities by simultaneously amplifying templates from Bacteria, Archaea, and Eukaryota in a single PCR reaction. Despite the potential to show all rRNA gene relative gene abundances, they are rarely used due to concerns about length bias against 18S amplicons and bioinformatic challenges converting mixed 16S/18S sequences into amplicon sequence variants. We thus developed 16S and 18S rRNA mock communities and a bioinformatic pipeline to validate this three-domain approach. To test for length biases, we mixed eukaryotic and prokaryotic mocks before PCR, and found consistent two-fold underestimation of longer 18S sequences due to sequencing but not PCR bias. Using these mocks, we show universal V4-V5 primers (515Y/926R) outperformed eukaryote-specific V4 primers in observed vs. expected abundance correlations and sequences with single mismatches to the primer were strongly underestimated (3-8 fold). A year of monthly time-series data from a protist-enriched 1.2-80 μm size fraction yielded an average of 9% 18S, 17% chloroplast 16S, and 74% prokaryote 16S rRNA gene amplicons. These data demonstrate the potential for universal primers to generate quantitative and comprehensive microbiome profiles, although gene copy and genome size variability should be considered - as for any quantitative genetic analysis.

## Introduction

Microbial communities of unicellular organisms are dynamic, diverse communities made up of bacteria, archaea, eukaryotes that interact with one another and their environment. Studying all these components together is essential for understand how the ecosystem functions as a whole (Fuhrman et al., 2015; Needham et al., 2018; Chénard et al., 2019), though single components are most typically studied alone due in part to the perception that separate assays are required for each. Since high-throughput DNA sequencing was introduced, SSU rRNA sequencing has been widely used for analyzing microbial community structure - especially for prokaryotes by targeting the 16S rRNA gene (Sogin et al., 2006). Analyses focusing on eukaryotic communities with 18S rRNA sequencing, however, are not as common partly because hypervariable regions are mostly longer than early sequencing lengths (Amaral-Zettler et al., 2009). Until recently, with current sequencing capacities, two regions, V4 and V7-9, have become commonly used for planktonic eukaryotic community profiles (Amaral-Zettler et al., 2009; Stoeck et al., 2010; Balzano et al., 2015; De Vargas et al., 2015). To study metazoan host-dominated communities, however, a concern is that universal 16S&18S primers would mainly amplify 18S host sequences, overwhelming the 16S and non-host 18S microbiome, and thus blocking primers or universal non-metazoan primers are required in these studies (Del Campo et al., 2019).

Despite these methodological developments, the question of how well the entire sequencing and analysis pipeline recovers the true abundance of rRNA genes found in the natural community has received less attention. In pelagic marine environments, studies have underscored the importance of careful primer design for accurately resolving natural communities, e.g. correcting the severe underestimate of the SAR11 clade that occurred with one of the most popular primers (Apprill et al., 2015; Parada et al., 2016). In addition, validation and inter-comparison of primer performance has also been facilitated by the development and application of microbial internal standards or “mock communities” to PCR amplicon analysis (hereafter “mocks”). The application of mocks to the PCR amplification, sequencing, and analysis protocol has demonstrated that even well-designed primers (515Y/926R vs. 515Y/806R) differ in terms of their ability to quantitatively recover natural abundance patterns (Parada et al., 2016). Including mocks in a sequencing run can also verify instrument performance, thus avoiding improper ecological conclusions. For example, mocks revealed that an unknown technical issue affecting a single sequencing run inexplicably caused an entire major taxon to be missing in output data and altered abundances of other taxa (Yeh et al., 2018). More recently, it has been shown that amplicon methods can be made even more quantitative by the addition of internal DNA standards (i.e. added to samples before extraction and purification of DNA). This allows normalization of amplicon data closer to true abundances found in seawater (except for lysis efficiency variations) and was found to be consistent with other, extensively-validated methods (Lin et al., 2019).

Bioinformatic methods used for amplicon sequence analysis have also evolved considerably, with initial efforts focusing on how well algorithms resolve true biological sequences by clustering sequences into operational taxonomic units (OTUs) at a certain similarity threshold. This effort has culminated in the development of “denoising” algorithms that are designed to recover true underlying biological sequences to the individual base (i.e. amplicon sequence variants; ASVs) by endeavoring to eliminate sequencing and PCR errors (Eren et al., 2015; Callahan et al., 2016; Amir et al., 2017). Unlike OTU clustering that must analyze sequences all together into often vaguely-defined or “fuzzy” units that change study-by-study, denoising methods aim to better account for batch effects across multiple sequencing runs, and are able to analyze sequences either sample-by-sample (Deblur) or run-by-run (DADA2), which greatly reduces computational demand (Callahan et al., 2016).

Collectively, these studies show how PCR amplicons can generate quantitative data that allows microbial community composition to be measured alongside other oceanographic variables. However, choosing an appropriate sequencing strategy remains a significant challenge given the diverse primers and sequencing technologies currently available. In order to maximize overall utility, it is highly desirable to keep costs low while generating data with high phylogenetic resolution. Parada et al. (2016) have previously described a universal primer set (515Y/926R) that simultaneously amplifies 16S and 18S rRNA in a single PCR reaction. Because of their universal nature, these primers measure both eukaryotic microbes and prokaryotes and can provide insights into processes such as predation, parasitism, and mutualism (Needham and Fuhrman, 2016; Needham et al., 2018).

However, analyzing data generated from universal 515Y/926R primer set has several potential challenges First, mixed 16S and 18S amplicon sequences present bioinformatic challenges since the two types of amplicons must be analyzed differently. This is because with current Illumina technology, the forward and reverse reads do not overlap for the 18S amplicon (575-595 bp), meaning that they cannot be merged as is typical for 16S amplicon analysis. Second, PCR and sequencing both discriminate against longer amplicons (Kittelmann et al., 2013), yet we lack a quantitative understanding of PCR and sequencing biases against longer 18S amplicons. These biases can potentially be detected via mock community analysis, specifically collections of known 16S or 18S rRNA gene fragments (Bradley et al., 2016; Parada et al., 2016). Yet to our knowledge, there have not been tests with mixed mock communities consisting of both 16S and 18S rRNA genes.

In this study, we present results from mock communities designed to validate the 515Y/926R primer set with particular emphasis on its performance with 18S sequences in comparison to commonly-used 18S-specific primer sets. We also present a bioinformatics workflow designed for mixed 16S and 18S amplicons that generates ASVs differing by as little as a single base, and reproducibly recovers the known exact sequences from the mock communities. This workflow, which uses common tools such as cutadapt (Martin, 2011), bbtools (http://sourceforge.net/projects/bbmap/), DADA2 (Callahan et al., 2016), deblur (Amir et al., 2017) and QIIME 2 (Bolyen et al., 2018), simplifies sequence analysis for mixed 16S and 18S amplicons, and allowed us to rigorously test the performance of two different denoisers (DADA2 and deblur) with a variety of different data types. We also rigorously examined biases between 16S and 18S amplicons at the PCR and sequencing steps. Lastly, we analyzed natural marine samples collected from San Pedro Ocean Time-series (SPOT) using the same workflow to examine the application of 515Y/926R to environmental samples.

## Results

### Effects of trim length on 18S denoising

Since 18S amplicons amplified with 515Y/926R are too long (∼575-595 bp) for forward and reverse reads to overlap (even with 2 × 300bp sequencing), we decided to trim reads to fixed lengths before denoising. Trimmed reads are concatenated either before (q2-dada2 and q2-deblur QIIME 2 plugins) or after denoising (standalone DADA2 R package). As quality profiles of reverse reads vary widely among runs, a trim length which worked equally well across runs needed to be determined. We therefore systematically decreased the trim length of reverse reads from 220 to 100 bp while fixing trim length of forward reads at 220 bp. Three criteria were then used to compare denoiser performance; 1) percent reads that perfectly matched *in silico* sequences, 2) R-squared values obtained by plotting the expected abundance of staggered mock community against the sequenced staggered mock community on a log (x+0.001) scale, and 3) percent reads removed after denoising (Table 1). Deblur successfully recovered staggered mock communities in the proportions expected regardless which trim length was chosen, and we did not observe any sequences without an exact match to the known reference sequences. DADA2, however, produced up to 0.5 % of reads that did not perfectly match the mock communities, though it performed slightly better when concatenating reads after denoising (0.3 % of reads did not perfectly match *in silico* sequences). By blasting these reads against *in silico* sequences, we found that these reads could be accounted for by sample bleed-through or contamination as they had more than 1-mismatch to the *in silico* sequences, which were less likely produced by PCR/sequencing errors. Although deblur never produced such putative erroneous ASVs, it removed a large fraction of reads (∼75%) compared with DADA2 (∼25%), yielding fewer observations in the final ASV table (Table 1). According to the three criteria defined above, denoiser performance was relatively consistent among runs at a trim length for the reverse read of 200 bp. Therefore, this length was used for the rest of the analysis.

**Table 1.** Effects of trim length on 18S staggered mock community.

| Trim<br>length<br>of<br>reverse<br>reads<br>(bp) | Deblur |  |  | DADA2 R package |  |  | DADA2 QIIME 2 version |  |  |
| --- | --- | --- | --- | --- | --- | --- | --- | --- | --- |
| | Percent<br>reads<br>perfectly<br>match in<br>silico | $r^2$<br>expected<br>vs.<br>observed | Percent<br>reads<br>removed<br>after<br>denoising | Percent<br>reads<br>perfectly<br>match in<br>silico | $r^2$<br>expected<br>vs.<br>observed | Percent<br>reads<br>removed<br>after<br>denoising | Percent<br>reads<br>perfectly<br>match in<br>silico | $r^2$<br>expected<br>vs.<br>observed | Percent<br>reads<br>removed<br>after<br>denoising |
| 220 | 100 | 0.77 | 73.5 | 99.5 | 0.76 | 35.8 | 99.4 | 0.77 | 20.0 |
| 210 | 100 | 0.77 | 72.6 | 99.6 | 0.74 | 31.2 | 99.4 | 0.77 | 29.3 |
| 200 | 100 | 0.77 | 72.3 | 99.7 | 0.76 | 28.3 | 99.5 | 0.77 | 27.3 |
| 190 | 100 | 0.77 | 71.2 | 99.8 | 0.76 | 25.8 | 99.6 | 0.77 | 25.1 |
| 180 | 100 | 0.77 | 70.2 | 99.8 | 0.76 | 24.3 | 99.6 | 0.77 | 23.4 |
| 170 | 100 | 0.76 | 71.5 | 99.9 | 0.76 | 23.3 | 99.7 | 0.76 | 22.4 |
| 160 | 100 | 0.76 | 70.4 | 99.9 | 0.76 | 22.5 | 99.6 | 0.76 | 21.6 |
| 150 | 100 | 0.76 | 69.3 | 99.9 | 0.76 | 21.7 | 99.7 | 0.76 | 20.7 |
| 140 | 100 | 0.76 | 70.2 | 99.9 | 0.76 | 20.9 | 99.8 | 0.76 | 19.5 |
| 130 | 100 | 0.76 | 69.2 | 100.0 | 0.76 | 20.4 | 99.9 | 0.76 | 18.9 |
| 120 | 100 | 0.76 | 68.3 | 100.0 | 0.76 | 19.8 | 99.8 | 0.76 | 18.1 |
| 110 | 100 | 0.76 | 67.4 | 100.0 | 0.76 | 19.3 | 99.8 | 0.76 | 17.6 |
| 100 | 100 | 0.76 | 66.4 | 100.0 | 0.76 | 18.8 | 99.9 | 0.76 | 16.9 |

### Comparison of 16S and 18S universal primers (515Y/926R) and eukaryote-specific primers (V4F/V4R and V4F/V4RB) with 18S mock communities

Our 18S mock communities are mixtures of a number of nearly full-length 18S rRNA genes with known concentrations, and were designed to represent the major eukaryotic groups found in marine environments, including Haptophyta, Dinophyta, Ochrophyta, Ciliophora, Cercozoa, Radiolaria, and Metazoa. Among them, Prymnesiales (Haptophyta) has a single mismatch to the reverse primer V4R (at the 3’ end), and three Dinophyta species (Lingulodinium, Dino-Group-II_b, and Gymnodinium) have a single mismatch to the reverse primer 926R. As the abundances of taxa in mock communities are known *a priori*, they can be used to test which primer set and denoising algorithm recover the community composition most closely to what is expected (Fig. 2).

**Figure 1.**
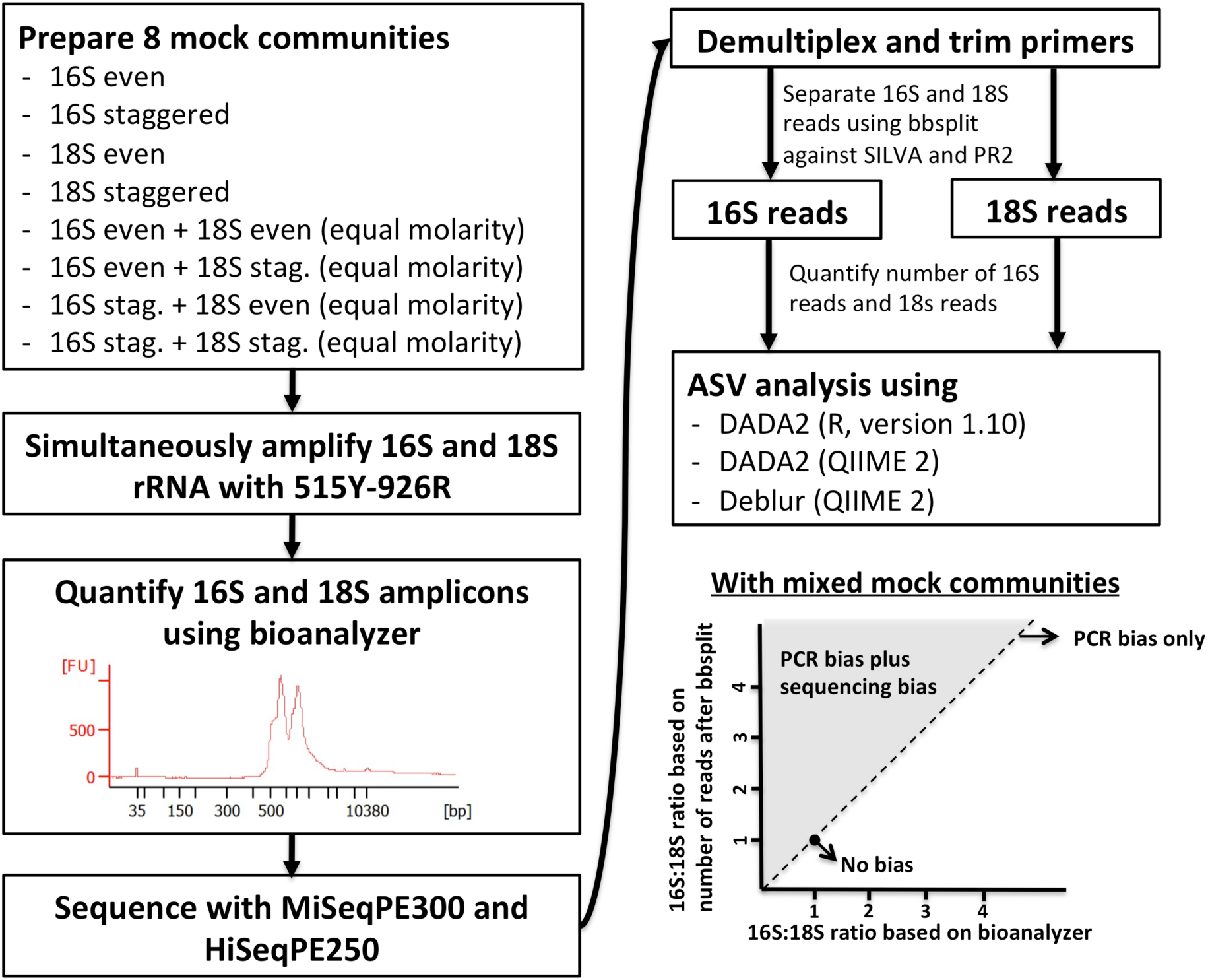
Experimental design. 8 mock communities were amplified using the 515Y/926R primers. The amplicons of mixed mock communities were analyzed using a Bioanalyzer to quantify the PCR bias against 18S amplicons. After sequencing, 16S and 18S reads were then separated through an *in-silico* sorting step and the number of 16S and 18S reads counted to quantify the sequencing bias against 18S. Hypothetically, if there is no bias, all the mixed mocks are located at a single point (1,1). If there is only PCR bias, all the data points will be at the one-to-one line. If there are PCR and sequencing bias, all the data points will be located above the one-to-one line (gray area). The slope indicates the sequencing bias.

**Figure 2.**
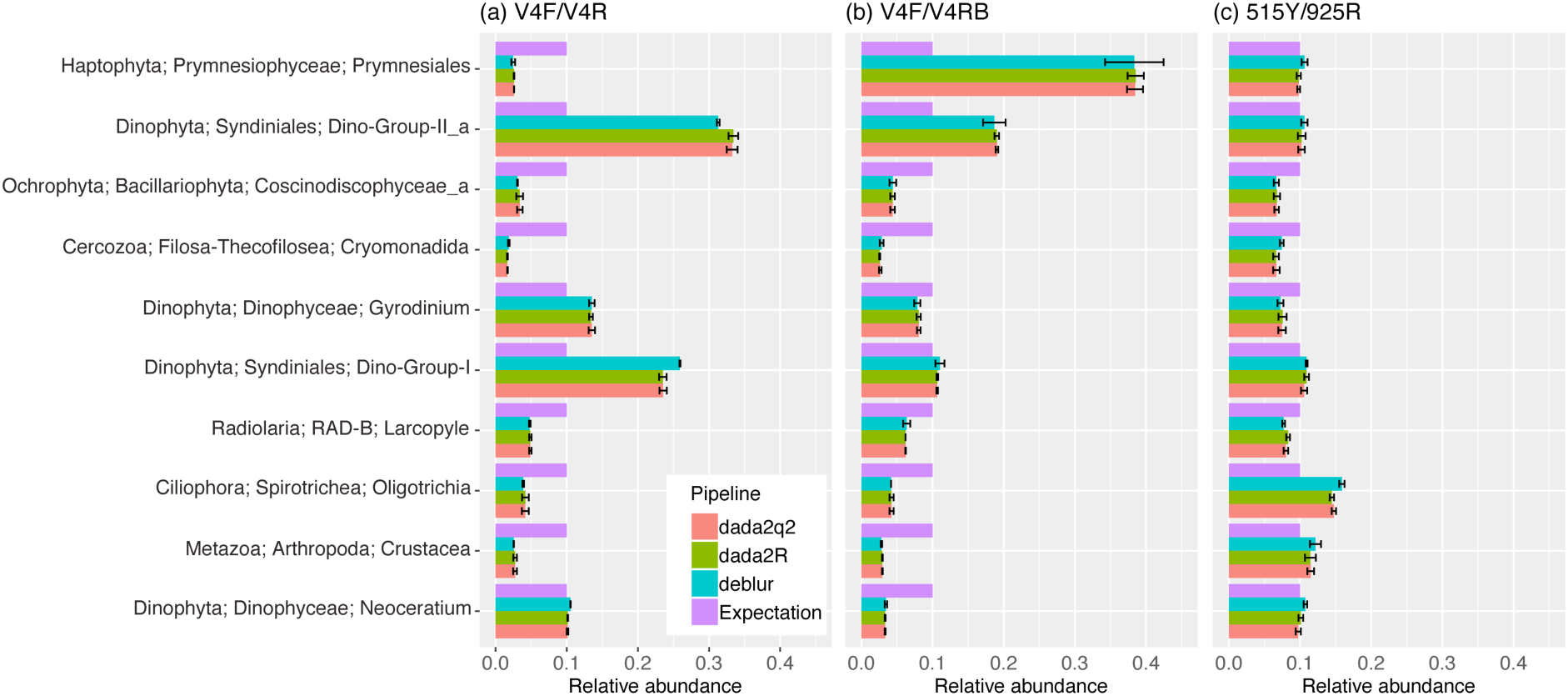
Comparison of even 18S mock communities amplified with V4F/V4R (a), V4F/V4RB (b), and 515Y/926R (c).

For 18S even mock communities, V4F/V4R underestimated Prymnesiales (Haptophyta) by ∼4-fold, presumably because of a single nucleotide mismatch on the 3’ end of the reverse primers (Fig. 2a) (Stoeck et al., 2010). On the other hand, the V4F/V4RB primers that do not have any mismatches overestimated Prymnesiales (Haptophyta) by ∼4-fold (Fig. 2b) (Balzano et al., 2015) while the 515Y/926R primers produced a community composition similar to that expected (Fig. 2c). The patterns were consistent among denoising pipelines (Fig. 2).

For 18S staggered mock communities, similar results were found. V4F/V4R underestimated Prymnesiales (Haptophyta) by ∼5-fold (Fig. 3a), and V4F/V4RB overestimated Prymnesiales (Haptophyta) by ∼3-fold (Fig. 3b). 515Y/926R underestimated three Dinophyta species (with single primer mismatches) to varying degrees (Lingulodinium, ∼8-fold; Dino-Group-II_b, ∼3-fold; Gymnodinium, ∼4-fold) (Fig. 3c). However, there was no relationship between degree of underestimation and locations of primer mismatch (Lingulodinium, −11 bases from the 3’ end; Dino-Group-II_b, −12 bases from the 3’ end; Gymnodinium, −2 bases from the 3’ end). The patterns were consistent among denoising pipelines (Fig. 3). Overall, the observed mock community composition was more similar to the expected with 515Y/926R (slope=0.88, r^2^=0.76), especially after removing three mismatched Dinophyta species (slope=0.97, r^2^=0.97), followed by V4F/V4RB (slope=0.79, r^2^=0.87) and V4F/V4R (slope=0.67, r^2^=0.65) (Fig. 4).

**Figure 3.**
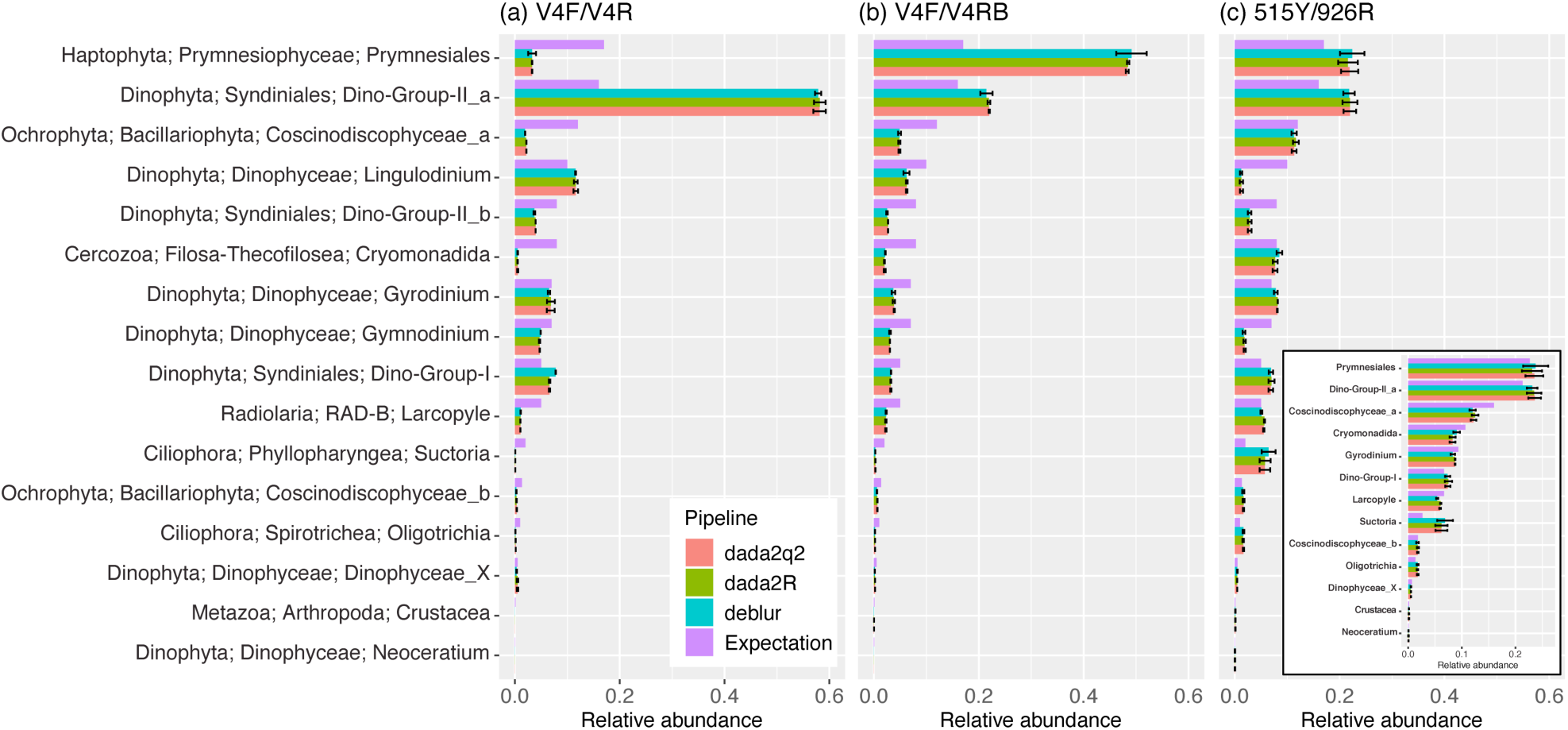
Comparison of staggered 18S mock communities amplified with V4F/V4R (b), V4F/V4RB (b), and 515Y/926R (c). The insert in (c) shows only taxa with perfect matches.

**Figure 4.**
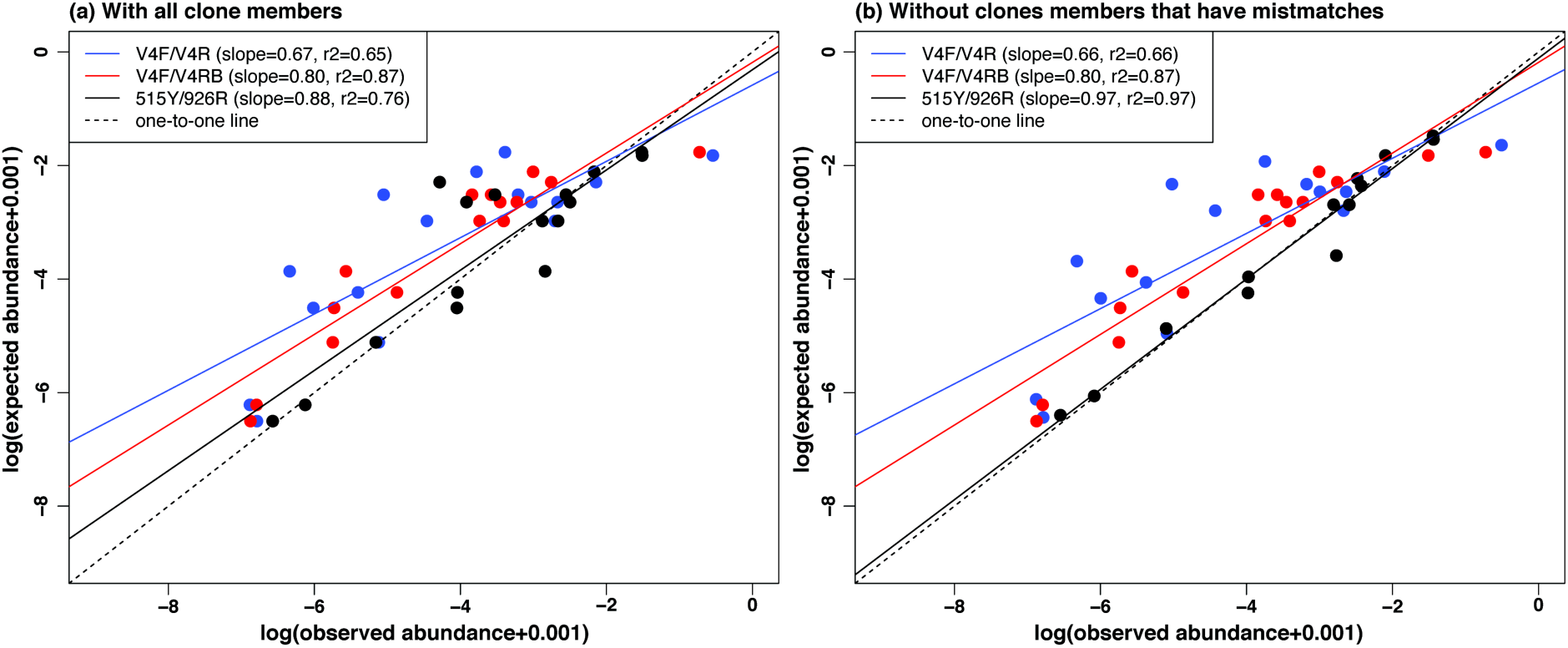
Expected staggered 18S mock communities plotted against observed staggered 18S mock communities amplified with different primers pairs (a) and without clone members that have mismatches on the given primer pairs (b).

### Estimation of PCR and sequencing bias against 18S reads in mixed mock communities

To test for length-based PCR bias against 18S reads, 18S mock communities were mixed with 16S mock communities in equimolar amounts prior to PCR amplification. The mixed mock communities were then PCR amplified, products analyzed on a Agilent 2100 Bioanalyzer, and then sequenced (Fig. 1). The bioanalyzer is a electrophoretic instrument that accounts for differences in sequence length in estimating the molarity (i.e., copies of DNA per unit volume) for DNA inputs. Based on bioanalyzer traces that separately quantify the abundance of 16S and 18S amplicons, there was little systematic PCR bias (about 0.7-1.3-fold) against 18S PCR products when using the 18S even mock communities that have no primer mismatches to 515Y/926R (Fig 5, orange and blue dots, x axis only). When the 18S staggered mocks were included (with three Dinophyta species that have one mismatch to the reverse primer, 926R), there was considerably more PCR bias, up to 3-fold (Fig 6 green and purple dots, x axis only). The mixed amplicons were then sequenced and split into 16S and 18S reads pools by an *in silico* sorting step. By comparing ratios in the bioanalyzer outputs and the raw read counts after *in silico* sorting, we observed that there was typically a 2-fold sequencing discrimination against 18S reads (Fig. 5), which is consistent regardless of community types (even, staggered) and sequencing runs.

**Figure 5.**
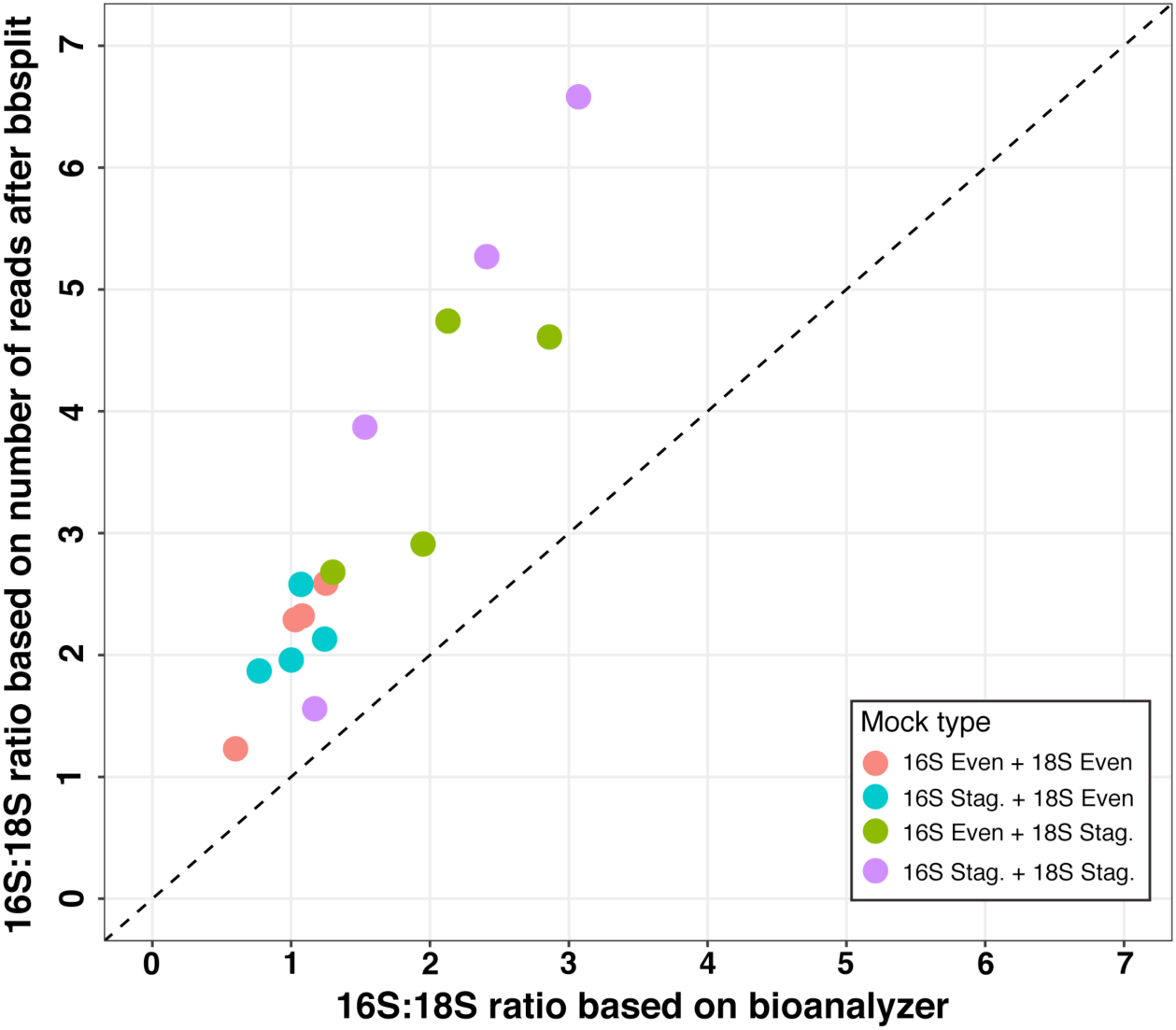
Comparison of PCR and sequencing biases among four mixed 16S and 18S mock communities, each combined in 1:1 molar ratios. The X axis shows ratios in the PCR products, and the Y axis shows the ratios in the final sequences, including biases from PCR plus sequencing. The data points all occur above the dashed 1:1 line, indicating most biases are from sequencing. Note for 18S even mocks (orange and blue) the PCR products have a bioanalyzer output ratio near 1, indicating little PCR bias. The staggered 18S mocks (green and purple) include 3 members with primer-template mismatches and correspondingly more PCR bias visible on the x axis. In all cases the final reads show about 2-fold more bias than the PCR biases alone.

**Figure 6.**
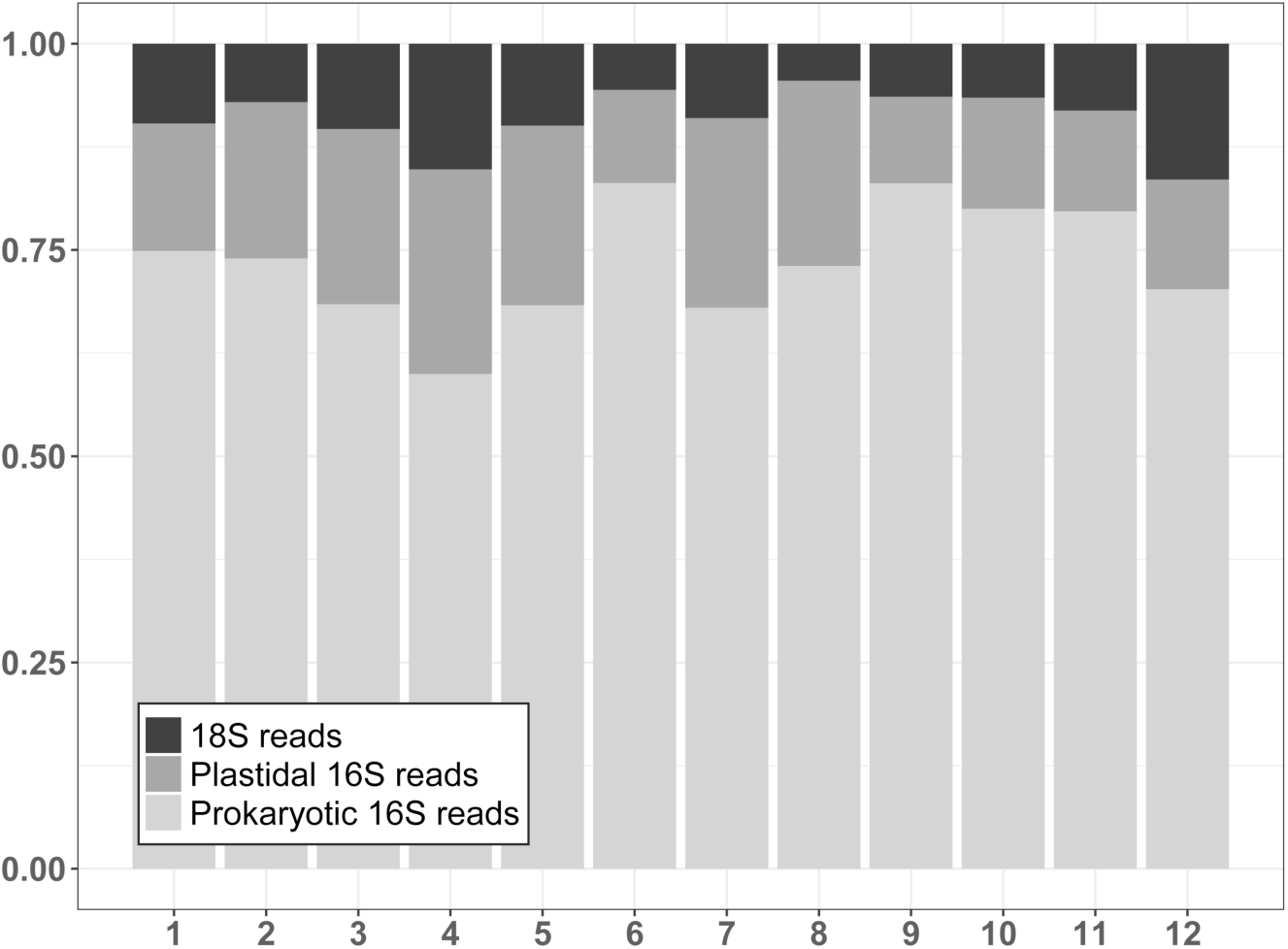
The composition of reads found in 1.2-80 μm size fraction of seawater samples collected from SPOT in 2014. Numbers on the X axis are months.

### The application of the 515Y/926R primer pair to field samples

To examine the application of universal primers (515Y/926R) to natural samples, surface seawater samples from a larger size fraction (1.2-80 μm) collected from the San Pedro Ocean Time-series (SPOT) location in 2014 were analyzed. On average, 9% of reads were 18S, 17% of reads were plastidal 16S reads, i.e. chloroplasts in photosynthetic eukaryotes (excluding dinoflagellates whose chloroplasts are not detected, Needham and Fuhrman (2016)), and 74% of reads were prokaryotic 16S (Fig. 6). A total of 2394 ASVs were identified across all samples (540 18S ASVs were affiliated to 228 orders; 442 plastidal 16S ASVs were affiliated to 81 orders; 1412 prokaryotic 16S ASVs were classified into 85 orders). Since Metazoa in this size fraction were patchy (maximum of 2% reads), mainly dominated by copepods Maxillopoda (Fig. 7a), they were separately analyzed in the community composition of 18S reads (Fig. 7b). 18S reads were dominated by Dinophyceae, Spirotrichea, Syndiniales, Ciliophora, Mamiellophyceae, MAST, Bacillariophyta, RAD-B, Prymnesiophyceae, and Polycystinea (Fig. 7b). For plastidal 16S reads, phytoplankton communities were dominated by Prymnesiophyceae, Mamiellophyceae, Pelagophyceae, Dictyochophyceae, Bacillariophyta, Cryptophyceae, Chrysophyceae-Synurophyceae, Raphidophyceae, Prasinophyceae, Bolidophyceae, and Chrysophyceae (Fig. 7c). The prokaryotic 16S reads showed that prokaryotic communities were mainly dominated by SAR11, Synechococcales (i.e. Cyanobacteria), Flavobacteriales, Rhodobacterales, Actinomarinales, Verrucomicrobiales, Puniceispirillales, Rhodospirillales, Opitutales, and Cellvibrionales. (Fig. 7d). The phytoplankton and prokaryotic communities had similar seasonal patterns, whereas non-phytoplankton eukaryotic communities were less predictable.

**Figure 7.**
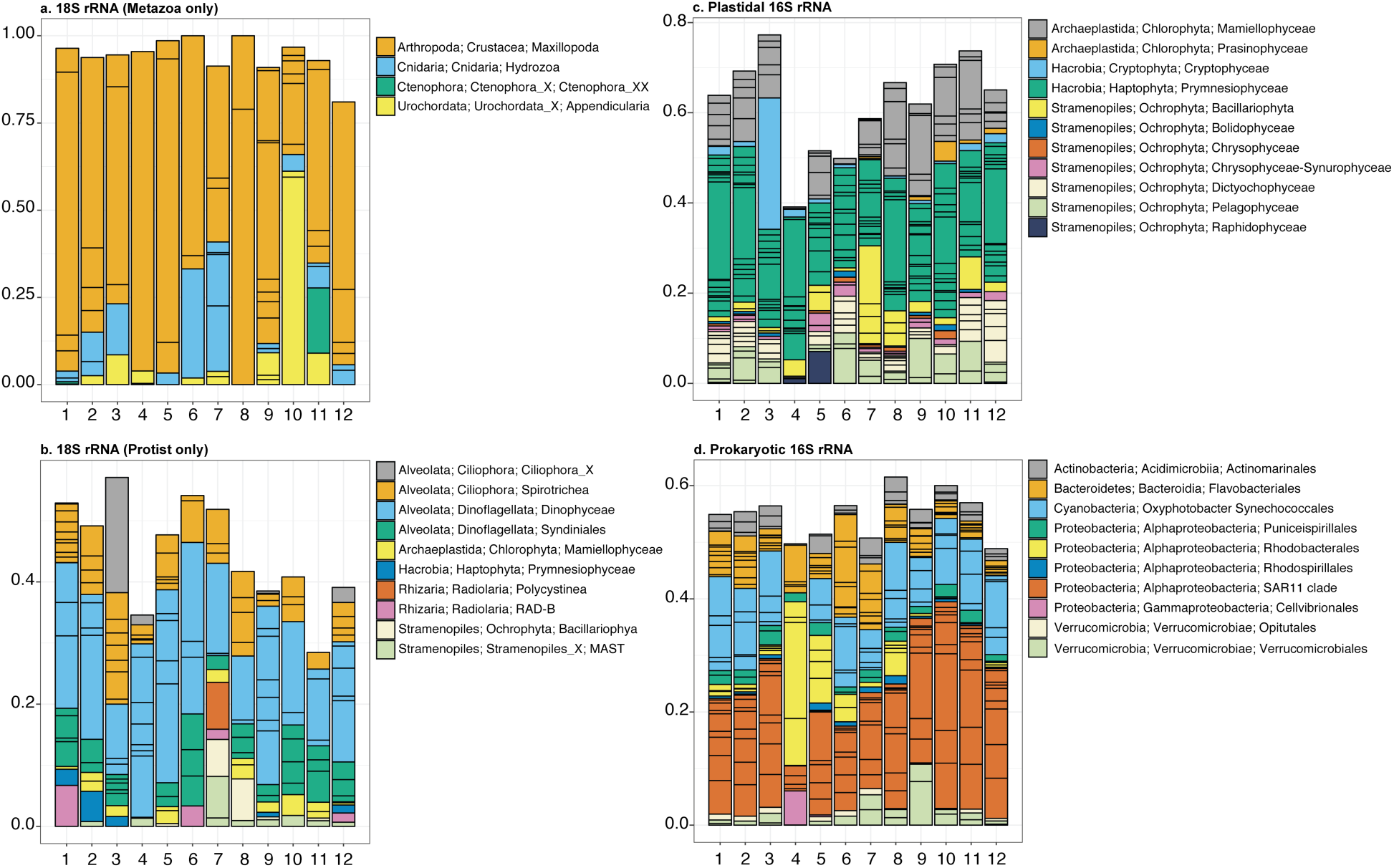
The monthly community composition of 1.2-80 μm size fraction of seawater samples collected from SPOT in 2014 at class level for eukaryotes and at order level for prokaryotes. Only the dominant ASVs (the average relative abundance is greater than 0.5%) were shown. Each box represents single ASV.

## Discussion

This study shows that the 3-domain universal primer (515Y/926R) can resolve community composition quantitatively for 16S and 18S rRNA in a single PCR reaction, with biases we could quantify and manage. We were able to investigate the biases relevant to the use of these primers in a natural setting through the use of 18S mock communities first applied here, separately and in concert with 16S mocks.

Unlike 16S rRNA sequencing, 18S rRNA sequencing using 515Y/926R is bioinformatically challenging because the amplicon is too long (∼575-595 bp) for forward and reverse reads to overlap with current Illumina sequencing capacities. A simplified solution in such a situation might be to use forward reads only instead of merged paired-end reads, because the quality of reverse reads is generally worse than forward reads, and errors near the 3’ end cannot be corrected without overlapping paired-end reads. However, using only forward reads sacrifices phylogenetic resolution. Therefore, acquiring paired-end information without producing extra artifacts becomes critical for 18S rRNA processing. To do so, we developed a bioinformatic workflow which allowed us to split mixed 16S/18S amplicons into two sequence pools by mapping reads to a curated 16S/18S reference database derived from SILVA and PR2. The workflow was validated with mixed mock communities, showing that the *in silico* sorting step is able to successfully separate 16S and 18S mocks apart without changing their composition. We analyzed 18S mock communities by trimming reads to fixed lengths before denoising. In this way, we not only removed error bases but also kept 18S ASVs comparable between runs. Our analysis showed that we were able to recover 18S mock communities as expected even when the trim length of reverse reads was reduced to 100 bp, implying that this analytical strategy can be used even for poor-quality sequences (as we sometimes see).

While analyzing the quantitative recovery of our mock communities and comparing 515Y/926R with other commonly-used 18S specific primers, we found a 3-8 fold underestimation when there was an internal primer mismatch, as was the case with three Dinophyta included in our 18S mock community with mismatches to the reverse primer 926R. The same issue was previously found with the original EMP primers (515C/806R, V4) that underestimated SAR11 by 8-fold (Apprill et al., 2015) and Thaumarchaea by 1.5-3 fold (Parada et al., 2016). Consistent with our findings, Bru et al. (2008) found that underestimation generally increased as mismatches were closer to the 3’ end of the primer, yet there was no predictable relationship between the position of mismatch and the degree of underestimation. The worst mismatches are at the 3’ end itself, as occurs with the V4R primer (Stoeck et al., 2010) for many common Haptophytes. This observation was the rationale for the creation of the V4RB primer with a 3’ degeneracy (Balzano et al., 2015) that greatly improves recovery of haptophytes that are often dominant in seawater (Berdjeb et al., 2018).

For sequences that matched the primers exactly, we found that the 515Y/926R primers quantitatively recovered the mock communities abundances for both 16S and 18S mocks (r^2^ = 0.97 for staggered mocks, observed vs. expected, Fig 3 and 4), indicating little preferential amplification of taxa that perfectly match primers. However, this did not apply to V4F/V4RB - although these primers perfectly matched all the clone members in the mock communities, they overestimated haptophytes by 3-4 fold. This discrepancy between results probably relates to methodological differences. V4RB has a considerably lower annealing temperature than V4F and a 2-step PCR is required (Stoeck et al., 2010). In contrast, amplification with the 515Y/926R primers can be done in a single step PCR reaction. This methodological difference may explain why the 515Y/926R primers more accurately recovered relative abundances of 18S taxa that have no mismatches versus the V4F/V4RB primers. These findings, together with the results of Parada et al. (2016), indicate that the 515Y/926R primers recover both 16S and 18S mock communities quantitatively when examined separately.

Since amplicon lengths of 16S and 18S rRNA gene are different, simultaneous amplification of 16S and 18S rRNA genes can lead to a length-based discrimination during PCR and sequencing stages, similar to what was previously reported to occur in general (Kittelmann et al., 2013). We endeavored to develop a quantitative understanding of the bias to better interpret results and to determine ideal sequencing depth for mixed communities. We therefore created mixed mock communities consisting of both 16S and 18S rRNA in equal molarity, amplified with 515Y/926R, and sequenced the resulting amplicons. The relative proportions of 16S and 18S amplicons were quantified using a bioanalyzer to evaluate PCR bias, and the number of recovered sequence reads were similarly counted (after splitting into 16S/18S pools in the bioinformatic pipeline) to determine the cumulative bias (both PCR and sequencing biases). We found very little PCR bias in even mocks that had no mismatches to the primers. As expected, the biases were 1.5-3-fold higher when samples included 18S staggered mock communities which contain three Dinophyta species that have one mismatch to the reverse primer. Moreover, a 2-fold sequencing bias (in addition to the aforementioned PCR biases) occurred in all combinations of mock community types. That suggests sequencing bias due to length differences is a consistent property of the Illumina sequencing platform, yet PCR bias due to primer mismatches is much less predictable. Thus, an evaluation of primer coverage across three domains, in actual field samples, may help better account for the PCR bias. Parada et al. (2016) found that 515Y/926R perfectly matches 86% of eukaryotes, 87.9% of bacteria, and 83.9% of archaea in the SILVA database, but we note that in actual practice the extent of mismatches in field samples depends on the particular taxa present and their proportions. We should also note that our 18S mock communities are very rich in alveolates such as dinoflagellates (3 of 10 in even, 7 of 16 in staggered) that tend to have mismatches to the 515F/926F primers; hence they probably overestimate the biases expected in most field samples.

Regarding pipeline recommendations, we found that all three pipelines (qiime2 q2-deblur plugin, qiime2 q2-dada2 plugin, and the standalone DADA2 R package) were capable of recovering our mock communities exactly, although we note several potential tradeoffs between the two algorithms we tested (deblur and DADA2) that should be considered. Both DADA2 and Deblur can accurately recover the mocks under most conditions. However, we found DADA2 sometimes generated 1-mismatches to reference sequences when challenged with noisy sequence data (data not shown) or sequencing runs where PCR amplification used inconsistent methodological parameters (e.g. differing PCR cycles or input concentrations). These results reinforce the advisability of running some sort of control to account for PCR errors or run-to-run variability (mocks or duplicate samples spread across runs, see also Yeh et al. (2018)).

Although our work shows that DADA2 has the potential to generate false positives, we also note potential drawbacks to deblur. While deblur never produced 1-mismatches to the mock reference sequences, it removed a much larger fraction of total input sequences (∼75%). This greatly reduces the sequencing yield, meaning that more sequencing is needed for the same coverage. In addition, the deblur algorithm differs significantly from DADA2 and there may be tradeoffs inherent in its design that are not apparent with the mocks. For example, deblur discards any reads deemed as errors whereas DADA2 attempts to correct sequencing errors (Callahan et al., 2016; Amir et al., 2017). Therefore, our evaluation with mock communities does not exclude the possibility that deblur may remove true biological variation deemed error sequences in natural samples where closely-related taxa coexist. Therefore, we recommend readers evaluate for themselves which denoiser is most appropriate for a given study, and consider the desired ultimate yield (of 16S and 18S sequences) when deciding sequencing depth.

To demonstrate the 515Y/926R full analytical pipeline with field samples, we analyzed monthly surface seawater samples collected from SPOT in 2014 from the 1.2-80 μm size fraction, which includes most protists, larger than average free-living prokaryotes and those attached to <80 μm particles. Even though this size fraction is enriched in protists (free living bacteria which are mostly < 1 μm are excluded), we found that 18S reads contributed an average of only to 9% of total reads, while chloroplasts averaged 17% and prokaryotes 74%. This proportion of 18S is similar to those reported from Southern California waters near SPOT (Needham and Fuhrman, 2016; Parada et al., 2016; Needham et al., 2018), but due to new results reported here we now know the 18S has a sequencing bias that underestimates in their copy number ∼2-fold. And while quantitative interpretation of 18S copy numbers is greatly complicated by the wide range of copies per genome (2 in some picoeukaryotes to >50,000 in some dinoflagellates and others, Zhu et al. (2005), the 16S of chloroplasts has a relatively smaller copy number variation and for phytoplankton the 16S plastid data probably more closely reflects relative biomass, e.g. chloroplast count, than does 18S (Needham and Fuhrman, 2016), though dinoflagellate chloroplasts are missed. The assay even detects the presence of multiple metazoa, in our case including copepods and larvaceans, the latter of which are voracious bacterivores. However, since these are relatively large individuals that may be patchily represented in samples, we bioinformatically separated them and recalculated proportions of protists alone (Fig. 7b).

Overall, this universal 515Y/926R primer pair is able to simultaneously examine the whole community structure across three domains, and we now have quantified the extent of sequencing biases that can be expected. This will allow us to better estimate cell abundances, biomasses, and overall inventories of all microbial (perhaps even some metazoa) taxa in a sample, and will facilitate improved interpretation of interactions, such as predation, parasitism, and mutualism. Although we now have quantified the biases associated with primer mismatch (e.g. the underestimation of some Dinophyta groups), the extent of underestimation is still unpredictable. Additionally, some protists groups (Diplonemea, Kinetoplastida, and Discosea) are known to have particularly long V4 regions (up to 800 bp) that are likely missed or seriously underestimated due to the amplification biases (R. Massana pers. Comm.). Thus, the potential limitations of interpretation of 18S results from this primer set, and probably any other primer set for a similar purpose, should be recognized when attempting to comprehensively analyze community composition.

## Materials and methods

### Mock community preparation

To generate even and staggered 16S mock communities, 11 and 27 clones of marine 16S rRNA genes were respectively prepared as previously described (Parada et al., 2016; Yeh et al., 2018). In the even 16S mock community, 11 clones were mixed with equal molarity. For the staggered 16S mock community, 27 clones were mixed with varying concentrations to roughly mimic marine prokaryote community composition observed at our sampling site.

To create even and staggered 18S mock communities, nearly full-length 18S rRNA clone libraries were prepared from the large size fraction (1.2-80 μm) of seawater samples collected from SPOT location. DNA was amplified using universal eukaryote primers Euk-A (5’-AACCTGGTTGATCCTGCCAGT-3’) and Euk-B (5’-GATCCTTCTGCAGGTTCACCTAC-3’) (Countway et al., 2005). 25-μl PCR mixtures contained 1.25X 5Prime Hot master mix (0.5 U Taq, 45mM KCl, 2.5 mM Mg2+, 200 μm dNTPs; Quanta Bio), 0.5 μM primers, and 1 ng of DNA. PCR conditions were performed as follows: an initial hot start step at 95°C for 2 min followed by 25 cycles of 95°C for 30 sec, 50°C for 30 sec and 68°C for 4 min. PCR products were then cloned using TOPO TA cloning kit with pCR4-TOPO Vector and One Shot Top 10 competent cells according to the manufacturer’s protocols. The cloned PCR products were sequenced using Sanger sequencing. Species identification was confirmed using BLASTn against the nt database. 16 clones were chosen to represent the major marine eukaryotic groups, including haptophytes, dinoflagellates, diatoms, ciliates, cercozoa, radiolaria, and copepods. Plasmids were purified using Qiagen plasmid plus 96 miniprep kit and then amplified with Euk-A and Euk-B using the same conditions described above. In the even 18S mock community, 10 clones were mixed with equal molarity. In the staggered 18S mock community, 16 clones were mixed with different concentrations. To mimic natural marine communities consisting of both eukaryotes and prokaryotes, 16S and 18S mock communities were mixed in four combinations (Fig. 1). Each mixed mock community was pooled with equal molarity after taking lengths into account (the average length of 16S mocks is 1425 bp, and the average length of 18S mocks is 1770 bp).

### Sample collection and DNA extraction

Samples were collected from 5m depth at San Pedro Ocean Time-series (SPOT) location in 2014. Approximately 12 L of seawater was sequentially filtered through 80-μm mesh, a 1.2-μm A/E filter (Pall, Port Washington, NY), and a 0.2-μm Durapore filter (ED Millipore, Billerica, MA). Filters were stored at −80°C until DNA extraction. This study only used DNA extracted from A/E filters (1.2-80 μm), which consist of both eukaryotic microbes and prokaryotes, for primer and pipeline testing purposes. DNA was extracted from A/E filters using a NaCl/CTAB bead-beating extraction protocol as described by Lie et al. (2013) with slight modification by adding an ethanol precipitation step after lysis to reduce the volume of crude extract, which helps minimize DNA loss during the subsequent purification.

### PCR and sequencing

To compare 16S/18S universal primers with eukaryote-specific primers, 18S mock communities were amplified with V4F (5’-CCAGCASCYGCGGTAATTCC-3’) and V4R (5’-ACTTTCGTTCTTGATYRA-3’), and V4F and V4RB (5’-ACTTTCGTTCTTGATYRR-3’) (Stoeck et al., 2010; Balzano et al., 2015). The only difference between these two primer pairs is the last nucleotide on the 3’ end of the reverse primer (A to R), which makes V4F/V4RB amplify haptophytes better (Balzano et al., 2015). Due to the considerably lower annealing temperature of the reverse primer, the full primers with indices and Illumina adaptors did not result in any PCR bands. Thus, 2-step PCR was required to obtain efficient amplification as is standard practice for this primer set (Stoeck et al., 2010; Balzano et al., 2015; Mahé et al., 2015; Pasulka et al., 2016). The first PCR mixtures contained 1X Phusion HF buffer (1.5 mM MgCl2), 300 μM dNTPs, 0.5 μM primers, 3% DMSO, 0.5 U Phusion High-Fidelity DNA polymerase (New England BioLabs Inc.), and 1 pg pure mock community. PCR cycles were as follows: 98°C for 1 min, 10 cycles of 98°C for 30 sec, 53°C for 30 sec, 72°C for 30 sec; and then 15 cycles of 98°C for 30 sec, 48°C for 30 sec, 72°C for 30 sec, and a final extension step of 72°C for 10 min. The second PCR reaction was performed with full primers that had barcoded indices and Illumina adaptors. PCR mixtures contained 1X Phusion HF buffer (1.5 mM MgCl2), 300 μM dNTPs, 0.5 μM primers, 3% DMSO, 0.5 U Phusion High-Fidelity DNA polymerase (New England BioLabs Inc.), and 2 μl of the PCR products from the first step. PCR cycles were as follows: 98°C for 1 min, 10 cycles of 98°C for 30 sec, 48°C for 30 sec, 72°C for 30 sec, and a final extension step of 72°C for 10 min.

With 16S/18S universal primers (515Y, 5’-GTGYCAGCMGCCGCGGTAA-3’; 926R, 5’-CCGYCAATTYMTTTRAGTTT-3’), as shown in Fig. 1, 8 different types of mock communities were amplified using single step PCR with full-length primers that had barcoded indices and Illumina adaptors. 25-μl PCR mixtures contained 1.25X 5Prime Hot master mix (0.5 U Taq, 45mM KCl, 2.5 mM Mg2+, 200 μm dNTPs; Quanta Bio), 0.3 μM primers, and 1 pg of pure mock community. PCR conditions were as follows: 95°C for 2 min, 30 cycles of 95°C for 45 sec, 50°C for 45 sec and 68°C for 90 sec, and a final extension step of 68°C for 5 min. Environmental samples were amplified using the same condition described above but with 0.5 ng of DNA. PCR products were cleaned using 0.8X Ampure XP magnetic beads (Beckman Coulter). Purified PCR products were quantified with PicoGreen and sequenced on Illumina HiSeq 2500 in PE250 mode and MiSeq PE300. For each sequencing run, multiple blanks (i.e. PCR negative controls) were included as internal controls, meaning PCR water was amplified, cleaned and sequenced as environmental samples with the same conditions described above. After sequence processing, blanks were used to check for contamination that comes from sample bleed-through due to “index hopping”.

### Sequence demultiplexing

Sequences were demultiplexed by reverse indices allowing no mismatches using QIIME 1.9.1 split_libraries_fastq.py (Caporaso et al., 2010). Then, forward barcodes were extracted using QIIME 1.9.1 extract_barcoded.py. The sequences were demultiplexed by forward barcodes allowing no mismatches using QIIME 1.9.1 split_libraries_fastq.py. The fully demultiplexed forward and reverse sequences were then split into per-sample fastq files using QIIME 1.9.1 split_sequence_file_on_sample_ids.py. The per-sample fastq files have been submitted to the EMBL database under accession number PRJEB35673.

### *In silico* processing of amplicons

Scripts necessary to reproduce the following analysis are available at github.com/jcmcnch/eASV-pipeline-for-515Y-926R. Demultiplexed amplicon sequences were trimmed with cutadapt, discarding any sequence pairs not containing the forward or reverse primer. We allowed an error rate of up to 20% to retain amplicons with mismatches to the primer. Mixed amplicon sequences were then split into 16S and 18S pools using bbsplit.sh from the bbtools package (http://sourceforge.net/projects/bbmap/) against curated 16S/18S databases derived from SILVA 132 (Quast et al., 2013) and PR2 (Guillou et al., 2013). The splitting databases used are available at https://osf.io/e65rs/. The two amplicon categories were then analyzed in parallel using qiime2 (Bolyen et al., 2019) or DADA2 implemented as the standalone R package (Callahan et al., 2016) as described below.

### 16S processing

16S sequences were analyzed using the DADA2 R package (Callahan et al., 2016), the QIIME 2 q2-dada2 plugin, and the QIIME 2 q2-deblur plugin (Amir et al., 2017) to compare different denoising outputs. We ran DADA2 in both R and qiime2 platforms to compare version differences (standalone R DADA2 = v1.10.1; qiime2-2018.8). With DADA2, forward and reverse reads were trimmed and filtered after inspecting their quality profiles. Filtered reads were used to make parametric error models for forward and reverse reads independently. Then, filtered reads were denoised based on run-specific error models. Denoised reads were then merged and remove chimeric reads. Note that the DADA2 R package allows us to exclude blanks from error model training, but qiime2 q2-dada2 plugin does not have this capacity. With deblur, reads were merged with qiime2 VSEARCH and filtered using qiime2 q-score-joined plugin. Filtered reads were then processed through the qiime2 q2-deblur plugin. 16S ASVs were classified with qiime2 classify-sklearn plugin against the SILVA 132 database subsetted to the amplicon region. 16S ASVs classified as Chloroplast were extracted based on the SILVA classifications and subsequently reclassified against the PhytoRef database.

### 18S processing

18S reads amplified with 515Y/926R are 575-595 bp, which is too long for forward and reverse reads to overlap, so we chose to trim reads to fixed lengths before the denoising step. While inspecting quality profiles, reverse reads were generally lower quality than forward reads and quality profiles varied among sequencing runs. To find trim lengths which worked equally well among runs, forward reads were trimmed to 220 bp and reverse reads were trimmed to varying lengths (100-220 bp). With the DADA2 R package, trimmed forward and reverse reads were used to make parametric error models independently. Then, trimmed reads were denoised based on run-specific error models. Denoised reads were concatenated and chimeric reads were removed. With the QIIME 2 q2-dada2 and q2-deblur plugins (which did not have an option for independent denoising and subsequent merging at the time of writing), forward and reverse reads were trimmed using bbduk.sh and concatenated using fuse.sh from bbtools package (http://sourceforge.net/projects/bbmap/). Concatenated reads were then processed as artificial single-end reads and chimeras were removed using QIIME 2 q2-dada2 and q2-deblur plugins. 18S ASVs were classified with qiime2 classify-sklearn plugin against both SILVA 132 and PR2 databases. Once processed into exact amplicon sequence variants (ASVs), 18S sequences were split into paired read files to indicate their non-overlapping nature.

### Validation of ASV algorithms by analysis of mock communities

ASVs were Blastn against *in silico* mock sequences to determine ASVs that perfectly matched *in silico* sequences, artifacts (1-mismatch to the *in silico* sequences), or contamination (more than 1-mismatch to the *in silico* sequences). To compare the performance of each pipeline, we determined 1) the percent of reads that perfectly matched *in silico* sequences, 2) the percent of reads removed after denoising, and 3) the R-squared values of linear regression between observed and expected abundances on a log (x+0.001) scale. The processing scripts in this study are available on Figshare at https://doi.org/10.6084/m9.figshare.11320388

### PCR and sequencing bias estimation

16S and 18S mixed mock communities amplified with 515Y/926R were run on a Agilent 2100 Bioanalyzer to quantify concentrations of 16S and 18S PCR products in each mixed mock community. Amplicons were analyzed with the High-sensitivity DNA assay kit according to the manufacturer’s instructions. Due to the length differences between 16S and 18S amplicons, the concentration of each amplicons were measured by checking peak area on Agilent 2100 Bioanalyzer using manual integration without altering the instrument-determined baseline. The 16S:18S ratio of molarity was used to determine PCR bias. Sequence pre-processing (i.e. bbsplit.sh) split reads into 16S and 18S pools. The 16S:18S ratio of the number reads was used to determine sequencing and PCR bias. The slope of the line derived from plotting the 16S:18S ratio from Bioanalyzer traces against 16S:18S ratio based on the number reads after the bbsplit step was used to define sequencing bias.

## Acknowledgments

We thank Mike Lee for selflessly taking the time to think and blog about how to split 16S from 18S informatically, which inspired the bbsplit method in our qiime2 workflow. This work was supported by NSF OCE 1737409, Gordon and Betty Moore Foundation Marine Microbiology Initiative grant 3779, and Simons Foundation Collaboration on Computational Biogeochemical Modeling of Marine Ecosystems (CBIOMES) grant 549943.

